# Effects of varenicline and cognitive bias modification on neural response to smoking-related cues: a randomised control trial

**DOI:** 10.1101/480566

**Authors:** Angela S. Attwood, Tim M. Williams, F. Joseph McClernon, Rachel Kozink, Sally Adams, Karl Scheeres, Amy Green, David Christmas, Marcus R. Munafò

**Author notes:** Corresponding author: Angela S. Attwood PhD, School of Psychological Science, University of Bristol, 12a Priory Road, Bristol BS8 1TU, United Kingdom.; E. Author email addresses: Tim Williams -, F. Joseph McClernon -, Rachel Kozink –, Sally Adams -, Karl Scheeres –, Amy Green –, David Christmas –, Marcus R. Munafò -. **Trial Registration:** Current Controlled Trials ISRCTN65690030. Date of registration: 41 30 January 2014.

## Abstract

Drug-related cognitive biases have been positively associated with drug-craving and increased likelihood of relapse. Cognitive bias modification paradigms have been developed to attenuate cognitive biases but there have been few studies that examined neural responses to these paradigms. This study compared neural responses following CBM and explored whether CBM effects were potentiated by varenicline administration. This was a double-blind placebo-controlled study with two between subject factors of drug (varenicline, placebo) and CBM (attend towards smoking cues, train away from smoking cues, control training) that recruited daily (≥ 10 cigarettes per day) non-treatment seeking smokers. Participants (n = 67, 53% female) were randomised to one-week of drug administration (varenicline or placebo) before attending a study session at which they were randomised to CBM condition, and underwent an fMRI scan were they were presented with smoking and neutral cues. Neural response to smoking (vs. neutral) cues, cognitive bias, craving and mood were assessed. There was no evidence of CBM effects on any outcomes. There was evidence of effects of varenicline on craving, with greater reductions in craving in the week preceding the study session in the varencline group (*p* = 0.04, *η*_*p*_^*2*^ = .06). There was also evidence of a drug by CBM interaction for neural responses (*z* = 3.78, *p* <0.001). Compared to placebo, varenicline was associated with greater activation in the right temporal middle gyrus in the CBM control condition, compared to an opposite effect in the CBM “attend towards” condition. These data suggest that CBM does not modify cognitive bias, subjective craving and mood, or neural response to smoking cues. There was also no evidence that CBM effects were potentiated by varenicline.

## Introduction

Smoking remains the leading cause of preventable death worldwide, with an estimated 6 million tobacco-related deaths occurring every year (World Health Organization, 2013). Despite many smokers reporting wanting to quit, few achieve long-term abstinence. This may be partly due to the presence of smoking-related cues in the environment, which through repeated and contingent pairing with drug administration, acquire powerful motivational properties that can precipitate craving and drug seeking (Foltin & Haney, 2000; Gray, LaRowe, & Upadhyaya, 2008; McClernon et al., 2016; Mucha, Pauli, & Angrilli, 1998; Muntaner et al., 1989).

Drug-related cognitive biases, characterised by selective or disproportionate attention allocation to drug cues, have been reported in users of a number of drugs and have been positively associated with drug craving (Field, Munafo, & Franken, 2009; Wakefield, Germain, & Henriksen, 2008), future drug use (Cox, Pothos, & Hosier, 2007), approach behaviours to drug-related cues (Franken, 2003) and increased likelihood of relapse (Marissen, Franken, Blanken, van den Brink, & Hendriks, 2007). Of particular importance, the drug-stimulus learning that is believed to underlie these biases is long-lasting, which makes an individual vulnerable to relapse long after initial cessation. In smokers, increased reactivity to smoking cues has been found to predict decreased likelihood of cessation (Abrams, Monti, Carey, Pinto, & Jacobus, 1988; Niaura, Abrams, Demuth, Pinto, & Monti, 1989; Waters et al., 2004). Consequently, reduction in cognitive bias is a potential target for therapeutic intervention.

There is evidence that it is possible to reduce cognitive biases using computer-based cognitive bias modification (CBM) paradigms that “train” individuals to allocate attention away from disorder-relevant cues. CBM has been shown to reduce cognitive and has also been associated with reduction in other symptoms such as low mood (Baert, De Raedt, Schacht, & Koster, 2010). Attwood and colleagues (Attwood, O’Sullivan, Leonards, Mackintosh, & Munafo, 2008) reported decreased cognitive bias in a group of smokers following one session of stimulus-avoidance CBM using a modified dot probe task. Compared to a group who had been trained to attend to smoking cues, there was evidence that the avoid group also showed attenuated craving in response to in vivo smoking cues in a subsequent cue exposure test (male participants only). A subsequent study in tobacco smokers found similar decreases in cognitive bias following CBM, but did not observe generalisation of these effects of other relevant behaviours (e.g., cigarette craving) or novel (untrained) stimuli (Field, Duka, Tyler, & Schoenmakers, 2009). The weak effects may be due to the limited number of sessions used in laboratory-based studies and that multiple session approaches may be more efficacious than single session CBM (Lopes, Pires, & Bizarro, 2014; McHugh, Murray, Hearon, Calkins, & Otto, 2010; Unrod et al., 2014). Taken together, the research suggests CBM may have some clinical utility but the effects are compromised by small effect sizes and issues with generalisation of the training effects.

There has been growing interest in the development of combination drug-behavioural therapies, in which a drug is used to augment the outcomes of a behavioural intervention (Swerdlow, 2012). This may offer a solution to the low efficacy and reliability of CET effects, if a suitable pharmacological agent can be identified. The smoking cessation pharmacotherapy varenicline acts as a partial agonist of the α4β2 nicotinic acetylcholine receptor and aids cessation by reducing cigarette craving and withdrawal symptoms. However, it has also been associated with a reduction in cue-related craving in humans (Ray et al., 2013), particularly following chronic administration (Brandon et al., 2011), and reductions in cue-induced reinstatement of drug taking in animals (Le Foll et al., 2012; Wouda et al., 2011). Therefore, varenciline may be a useful adjunct to CBM, particularly as it is already licensed as a smoking cessation aid.

The current study replicated earlier work by examining the effects of CBM on behavioural measures of cognitive bias (visual dot probe and modified Stroop), and extended the work in two important ways. First, we examined whether 7-day pre-treatment of varenicline enhanced the effects of CET on smoking cue reactivity and attentional bias. Second, using fMRI, we examined the neural responses to smoking cues following treatment. Neuroimaging studies suggest that drug-related cognitive biases are the result of a failure of cognitive regulatory systems to increase control in the presence of salient cues that increase processing in the reward and emotional centres of the brain (e.g., striatum, amygdala) (Hester & Luijten, 2013). Therefore, these additional measures offer important insight into the mechanisms underlying the effects of CBM, particularly as computer-based measures of cognitive bias are known to lack reliability (Ataya et al., 2012; Field & Christiansen, 2012).

In this study, participants attended two sessions, approximately one-week apart. Participants were randomised to receive either 7-day treatment of varenicline or matched placebo, prior to completion of 1-hour of CBM training. For CBM training, participants were further randomised (stratified by drug group) to one of three conditions: 1) training towards smoking cues (attend), 2) training away from smoking cues (avoid), or 3) control training (control). After training, participants underwent a cue reactivity task during an fMRI scan. We hypothesised that participants in the CBM-avoid condition would show a decrease in cognitive bias and neural response to smoking-related cues compared to those in the CBM-attend and CBM-control conditions, and that changes in cognitive bias and neural response, would be greatest in individuals trained to avoid smoking-related cues *and* treated with varenicline.

## Methods

### Participants

Daily smokers (≥10 cigarettes or 15 roll ups/day who smoke within one hour of waking) were recruited from the staff and students at the University of Bristol and the general population through existing participant databases, posters, online flyers and word of mouth. Participants were required to be between 18 and 40 years of age, fluent in English and registered with a General Practitioner. Exclusion criteria were pregnancy, breast feeding or at risk of pregnancy (i.e., females not using adequate contraception), substance misuse, high alcohol consumption (>35 units/week if female or >50 units/week if male) or caffeine consumption (> 8 cups of caffeinated beverage per day), current or past psychiatric disorder, clinically significant abnormality (including cardiology risk factors), use of medication (participants were required to be 8 weeks clear of any prescribed medication), known hypersensitivity to varenicline, high blood pressure or heart rate (systolic/diastolic >140/90 mmHg or heart rate >90 bpm), uncorrected visual or auditory impairment, and any condition that would make MRI scanning unsafe (e.g., metallic implants) or intolerable. The study was approved by the National Research Ethics Service (London Brent Committee, reference: 11/LO/1726). All participants gave written informed consent and were reimbursed £70 at the end of the study.

### Design

This was a double-blind, placebo-controlled study that used a 2 × 3 between-subjects design, comprising one factor of drug (varenicline, placebo) and one factor of CBM group (attend, avoid, control). For the behavioural assessments of cognitive bias (dot probe, modified Stroop), there was an additional within-subjects factor of cue type (smoking, neutral).

### Drug administration

Following initial consent and screening on day 0, varenicline (or matched placebo) was prescribed by a medical doctor for one week. Participants were told to take one tablet (0.5 mg) daily on days 1 - 3, two tablets daily (total 1 mg) on days 4 - 6, and one tablet (0.5 mg) on day 7, consistent with standard dosing regimen for smoking cessation. Participants completed daily diaries detailing the time at which the tablets were taken and any side effects. Participants attended their second session on day 7 (i.e., their last drug day) and were asked to take the drug in the morning prior to their study session.

### Randomisation

Participants were randomly assigned to drug and CBM groups (stratified by gender), but equal numbers of participants per group were maintained. Drugs were supplied by Pfizer and shipped to University Hospitals Bristol Pharmacy who prepared two batches of 36 bottles (one for male and one for females). Within each group of 36 bottles, 18 bottles contained 10 varenicline tablets (0.5 mg each) and 18 bottles contained 10 tablets of matched placebo. Each bottle was given a numeric identifier that enabled study staff involved in data collection to be fully blinded to drug condition.

In addition, an experimental collaborator (who had no direct contact with the study participants) prepared a numeric code using random number assignment software to further randomise participants to CBM groups. Randomisation was stratified so that equal numbers of male and female participants (n = 12) were allocated across the six experimental cells (drug treatment [2] × CBM group [3]).

### Measures and materials

#### Materials

Stimuli for the CBM and the fMRI cue exposure task comprised full-colour 32 smoking-related pictures and 32 neutral pictures. Smoking-related cues consisted of full-colour pictures of people smoking. Control cues consisted of full-colour pictures of people engaged in everyday activities (e.g., talking on the telephone, writing). Equal numbers of females and males were represented in each category. The set of cues pictures is the same as used in previous imaging studies (McClernon, Kozink, Lutz, & Rose, 2009; McClernon, Kozink, & Rose, 2008). For the cognitive bias modification task, an additional 4 picture pairs, unrelated to smoking, were used in practice and buffer trials.

#### Cognitive Bias Modification

Participants were randomised to complete a modified visual dot probe task designed to induce a biased cognitive response away from (avoid: n = 24), or towards (attend: n = 24) smoking-related cues, or a control condition (control: n = 24). Each task version comprised 768 trials. Each trial began with a fixation cross (500 ms), before a picture pair (smoking image, neutral image) was presented on a computer screen. The picture pair stayed on screen for 500 ms and then was replaced by a probe (small square or circle) in a location previously occupied by one of the pictures. Participants were required to identity whether the probe was a square or circle by pressing designated keyboard keys.

The majority of trials (n = 512) were training trials, presented in four blocks, and the remainder of trials (n = 256) were test trials. Half of the test trials (n = 128) were presented prior to the training trials (baseline test), and half (n = 128) after the training trials, in order to assess the effect of the CBM on cognitive bias. In the test trials, the probe appeared with equal frequency in the location of the smoking-related or neutral picture. In the training trials, the probe appeared in the location of the neutral picture on 75% of trials in the avoid condition, or the smoking-related picture on 75% of trials in the attend condition, or with equal frequency in the location of neutral and smoking-related pictures in the control condition. The inter-trial interval jittered between 750 ms and 1,250 ms. The tasks were programmed and presented using EPrime version 2 software (Psychology Software Tools Inc., Pittsburgh PA), and total task time was approximately 50 minutes.

#### Cognitive Bias Generalisation Test (modified Stroop)

A pictorial version of the modified Stroop task was used to investigate the effect of dot-probe CBM on a different measure of cognitive bias. The task began with 16 practice trials followed by two experimental blocks, each comprising 8 buffer and 96 experimental trials (i.e., 208 trials in total). For each trial a picture was presented (smoking-related or neutral) centrally on screen. The picture was surrounded by a coloured border and the participant was required to identify the colour of the border (red, blue, yellow or green) using colour-marked keys on the keyboard.

#### Questionnaires

Questionnaire measures included the Eysenck Personality Questionnaire – Revised (EPQ-R) (Eysenck & Eysenck, 1991), the Questionnaire of Smoking Urges - Brief (QSU-Brief) (Tiffany & Drobes, 1991), the Minnesota Nicotine Withdrawal Scale (MNWS) (Hughes & Hatsukami, 1986) and visual analogue scales (VAS) of mood and cigarette craving.

#### fMRI Acquisition

An anatomical and a fMRI scan were performed on the test day (Day 7) with a Siemens Magnetom Skyra 3T scanner. BOLD images were acquired using an EPI sequence (36 slices, TR = 2,500 ms, TE = 30 ms, FOV = 19.2 cm, matrix = 64 × 64, flip angle = 90^°^, slice thickness = 3 mm, resulting in 3 mm^3^ isotropic voxels). A T1-weighted structural image was acquired using an MP-RAGE sequence with a 0.9 mm^3^ isotropic voxel size and 192 slices. During the fMRI cue exposure procedure, smoking-related and control cues were presented in a boxcar design with four blocks per category. Participants were required to make a button press on each stimulus presentation to confirm they had seen the image (this did not terminate viewing time). Each block lasted 40 s, during which time 8 cues were presented. Before and after each block, a crosshair was presented for 5 s. Participants were then asked to rate cigarette craving on an 8-point scale (“none at all” to “extreme”). The scale was presented for 10 s followed by a crosshair for another 10 s. Thus, the total interblock-interval was 25 s. The sequence of events was controlled using EPrime version 2 software (Psychology Software Tools Inc., Pittsburgh PA), and total task time was approximately 10 min.

### Procedure

Individuals who responded to study advertisements were sent the full information sheet and completed a telephone screening to assess basic eligibility. Eligible participants were then booked in for a screening and baseline assessment session (Day 0). At this session full written informed consent was taken by a trained researcher, and then the screening procedure was completed. This included measures of expired breath alcohol and carbon monoxide, height, weight, blood pressure and heart rate. A urine screen was performed to test for recent drug use (all) and pregnancy (females). A medical doctor then completed a general physical and psychiatric health assessment, and prescribed the study medication if appropriate. Then participants completed a baseline assessment of cognitive bias (modified Stroop), questionnaires assessing personality (EPQ-R), cigarette craving and withdrawal (QSU, MNWS) and mood (VASs), and a practice version of the task that they would completed during the fMRI scan at the second visit (Day 7).

Participants were then sent away with the study medication, medication packaging information and a drug diary (which they were required to complete and return at the next visit). The second session (test day) was then scheduled for approximately one week later. This session fell on day 7 of their drug regimen.

On the test day (Day 7), participants returned with their drug diaries and any untaken medication. Prior to the scan, they completed the Stroop task followed by a short visual dot probe task that measured baseline cognitive bias. Participants then completed one version of CBM (avoid, attend, control) per the study randomisation. The test version of the dot-probe task was run again immediately post-CBM in order to assess changes in cognitive bias. Following this, participants completed a 4-minute anatomical scan and then the cue-exposure test during a 15 minute fMRI scan. After scanning, participants completed the modified Stroop task and questionnaires (QSU, visual analogue scales) again. At the end of the test session, participants were offered smoking cessation literature, debriefed, and reimbursed.

### Data analysis

The protocol for this study was published in Trials in October 2014 (Attwood, Williams, Adams, McClernon, & Munafo, 2014).

#### Cognitive bias analyses (visual dot probe test of CBM and Stroop)

All data were examined for outliers (defined as scores three or more standard deviations above or below the group mean). Removals are noted in text. Data were assessed for normality and transformed using log 10 transformations (or square root where data include zero scores) where deviations from normality were observed.

Mean reaction time data were extracted for each of the four variables from the visual dot probe tests that were completed before and after CBM training (pre-training neutral, pre-training smoking, post-training neutral, post-training smoking). Cognitive bias scores were calculated by subtracting RTs to probes that replaced smoking-related pictures from RTs to probes that replaced neutral pictures, so that positive scores represent a bias towards smoking cues and negative scores represent a bias towards neutral cues. These bias scores were used to examine cognitive training effects in a 2 (pre-, post-CBM) × 2 (varenicline, placebo) × 3 (attend, avoid, control) mixed model ANOVA.

A similar procedure was applied to Stroop data (test of cognitive bias generalisation). Mean reaction times and errors were extracted for smoking and neutral images during tasks completed before and after CBM training. Bias scores were calculated for each variable and used in the same 3-way mixed model ANOVA (detailed above), with exception that the subtraction was reversed (i.e., neutral scores were subtracted from smoking scores). This was done for ease of interpretation as (unlike the visual dot probe) slower scores represent cognitive bias on the Stroop task. Therefore, for Stroop data presented here positive scores represent a cognitive bias to smoking cues.

#### Questionnaire analyses

Withdrawal (MNWS), craving (QSU) and mood (VAS) data were analysed in two time phase analyses using ANOVA. We first assessed tonic craving (QSU) defined as craving during drug treatment. This analysis used baseline data from sessions one and two in a 2 (pre-drug, post-drug treatment) × 2 (varenicline, placebo) mixed model ANOVA. We then examined craving and mood change across CBM (i.e., on test day) in a series of 2 (pre-, post-CBM) × 2 (varenicline, placebo) × 3 (attend, avoid, control) mixed ANOVAs.

#### fMRI analysis

BOLD signal pre-processing was conducted in FSL version 5.0.1 (Jenkison et al, 2012) to remove noise and artefacts. The first two volumes of each run were discarded to allow for T1 stabilization. All functional images were corrected for head motion using rigid-body transformations (MCFLIRT; Jenkinson et al, 2002) and acquisition timing. Additional pre-processing steps included spatial smoothing using an 8 mm FWHM Gaussian filter, high-pass filtering, and registration to standard space using FLIRT.

Each participant’s fMRI data was then entered into a first-level voxel-by-voxel analysis using the general linear model. Each cue block (smoke, control) was modelled as a boxcar function convolved with a double-γ hemodynamic response function that begins at the onset of the first cue in the block and ends at the end of the block (duration = 40 s). A smoking>control cue contrast image was created and input into a random effects analysis. A 2 (varenicline, placebo) × 3 (attend, avoid, control) mixed-model whole-brain ANOVA was used to examine smoking cue reactivity (smoking greater than control) between each group. Activation was evaluated within an *a priori* mask of anatomical brain regions identified in a meta-analysis of cue reactivity (Tang, Fellows, Small, & Dagher, 2012): nucleus accumbens, caudate, putamen, temporal gyrus, anterior cingulate gyrus, amygdala, insula, posterior cingulate cortex, inferior frontal gyrus, and angular gyrus. Resulting activations within the mask were considered significant at p<0.001 (uncorrected) with a minimum cluster extent threshold of 20 contiguous voxels. Smoking cue greater than control cue contrast images for each participant were input into random effects regression analyses examining relations between post-training cognitive bias scores and brain cue reactivity.

#### Sample size determination

The effects of cognitive bias on brain responses to smoking cues have not been evaluated in previous research. Data from our previous studies (Attwood et al., 2008) indicated a likely increase in cognitive bias index of 30 ms in the attend group, and a decrease of 30 ms in the avoid group. We assumed that the change in the control condition would be intermediate (i.e., 0 ms). Using these estimates, we calculated that we would achieve greater than 80% power to detect a linear effect across the three study groups on change in cognitive bias index with a total sample size of n = 30, at an alpha level of 0.05. Due to an additional factor of drug group (varenicline vs. placebo), the actual sample size we recruited was n = 72, with n = 12 per experimental group.

## Results

### Characteristics of participants

Four participants withdrew from the study (two from varenicline/control CBM condition, and two from placebo/avoid CBM condition) and therefore the final sample comprised 68 participants (53% female). Participants were aged between 18 and 39 years (*M* = 23, *SD* = 5) and smoked between 10 and 25 cigarettes per day (*M* = 15, *SD* = 3). Alcohol Use Disorders Identifier Test scores ranged between 5 and 24 (*M* = 13, *SD* = 4). EPQ-R scores ranged between 3 and 17 (*M* = 8, *SD* = 3) for psychoticism, 0 and 20 (*M* = 7, *SD* = 4) for neuroticism, and 10 and 23 (*M* = 18, *SD* = 3) for extraversion. See Table 1. for participant characteristics by drug and CBM condition.

**Table 1.**
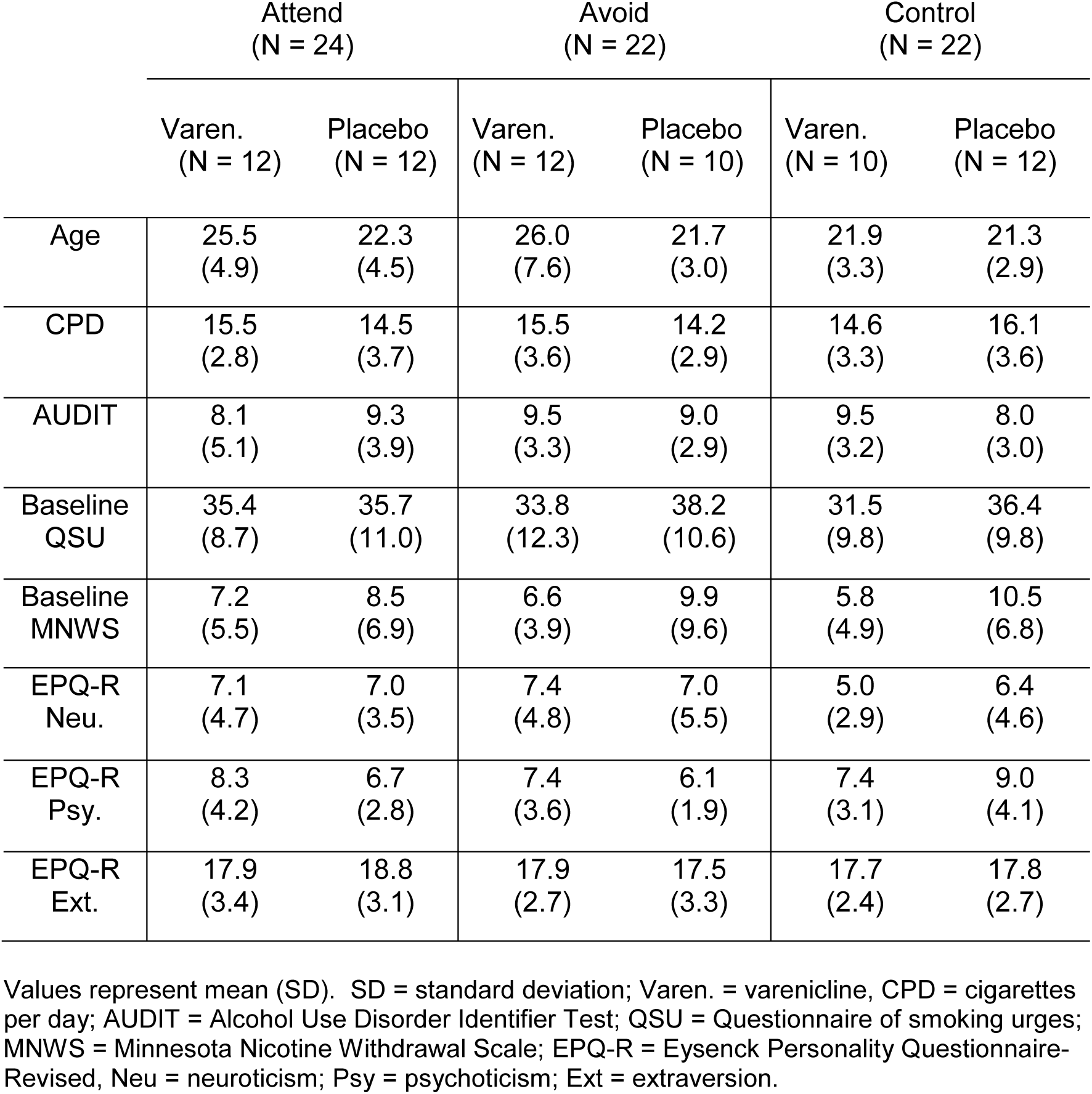
Mean (*SD*) participant characteristics, baseline craving and nicotine withdrawal

|  | Attend<br>(N = 24) |  | Avoid<br>(N = 22) |  | Control<br>(N = 22) |  |
| --- | --- | --- | --- | --- | --- | --- |
|  | Varen.<br>(N = 12) | Placebo<br>(N = 12) | Varen.<br>(N = 12) | Placebo<br>(N = 10) | Varen.<br>(N = 10) | Placebo<br>(N = 12) |
| Age | 25.5<br>(4.9) | 22.3<br>(4.5) | 26.0<br>(7.6) | 21.7<br>(3.0) | 21.9<br>(3.3) | 21.3<br>(2.9) |
| CPD | 15.5<br>(2.8) | 14.5<br>(3.7) | 15.5<br>(3.6) | 14.2<br>(2.9) | 14.6<br>(3.3) | 16.1<br>(3.6) |
| AUDIT | 8.1<br>(5.1) | 9.3<br>(3.9) | 9.5<br>(3.3) | 9.0<br>(2.9) | 9.5<br>(3.2) | 8.0<br>(3.0) |
| Baseline<br>QSU | 35.4<br>(8.7) | 35.7<br>(11.0) | 33.8<br>(12.3) | 38.2<br>(10.6) | 31.5<br>(9.8) | 36.4<br>(9.8) |
| Baseline<br>MNWS | 7.2<br>(5.5) | 8.5<br>(6.9) | 6.6<br>(3.9) | 9.9<br>(9.6) | 5.8<br>(4.9) | 10.5<br>(6.8) |
| EPQ-R<br>Neu. | 7.1<br>(4.7) | 7.0<br>(3.5) | 7.4<br>(4.8) | 7.0<br>(5.5) | 5.0<br>(2.9) | 6.4<br>(4.6) |
| EPQ-R<br>Psy. | 8.3<br>(4.2) | 6.7<br>(2.8) | 7.4<br>(3.6) | 6.1<br>(1.9) | 7.4<br>(3.1) | 9.0<br>(4.1) |
| EPQ-R<br>Ext. | 17.9<br>(3.4) | 18.8<br>(3.1) | 17.9<br>(2.7) | 17.5<br>(3.3) | 17.7<br>(2.4) | 17.8<br>(2.7) |

Values represent mean (SD). SD = standard deviation; Varen. = varenicline, CPD = cigarettes per day; AUDIT = Alcohol Use Disorder Identifier Test; QSU = Questionnaire of smoking urges; MNWS = Minnesota Nicotine Withdrawal Scale; EPQ-R = Eysenck Personality Questionnaire-Revised, Neu = neuroticism; Psy = psychoticism; Ext = extraversion.

### Cognitive bias modification test (visual dot probe)

Due to computer malfunction, post-training CBM data were not recorded for one participant, therefore post-training sample comprises 67 participants. This participant completed the allocated CBM training and therefore their data have been retained in all other analysis. Prior to the calculation of bias scores, reaction time data of the four primary variables (pre-training neutral, pre-training smoking, post-training neutral, post-training smoking) were assessed for normality, and there was evidence of positive skewness on three of the four variables. Therefore log 10 transformation was applied to the data prior to the bias calculations being performed. The 2 (pre-, post-CBM) × 2 (varenicline, placebo) × 3 (attend, avoid, control) mixed model ANOVA showed no main effects or interactions (*p*s >.23). These findings did not change if untransformed data were used (*p*s >.18).

### Cognitive bias generalisation test (modified Stroop)

For error data, three participants were identified as outliers in the pre-CBM condition and one participant was identified as an outlier in the post-CBM condition. These data were removed from main analysis. After data removal error data were not normally distributed and a square root transformation was applied to these data.

There was weak evidence of a drug × CBM interaction (*F* [1, 57] = 2.74, *p* = .073, *η*_*p*_^*2*^ = .09) for errors, reflecting a bias towards smoking (versus neutral) cues in the attend CBM condition (compared to avoid and neutral conditions) but only in individuals who had received placebo. In contrast, participants who received varenicline showed a bias towards neutral in the attend condition (compared to avoid and neutral conditions). The evidence for this effect was weaker when untransformed data were used (*p* = .18) and when outliers were included (*p* = .29). There was no evidence of any other main effects or interactions for Stroop error (*p*s >.13) or reaction time (*p*s >.15) data.

### Questionnaire data

#### Withdrawal across drug treatment

There was weak evidence of an effect of drug (*F* [1,66] = 3.34, *p* = .072, *η*_*p*_^*2*^ = .05) with lower nicotine withdrawal in the drug (*M* = 7.0, *SD* = 6.3) compared to placebo group (*M* = 9.8, *SD* = 6.3). There was no clear evidence of an effect of time or time by drug interaction (*p*s > .35).

#### Tonic craving across drug treatment

For QSU data, there was evidence of effects of time (*F* [1,65] = 33.61, *p* < .001, *η_p_^2^* = .34) and drug (*F* [1,65] = 6.06, *p* = .017, *η_p_^2^* = .09), which were subsumed under a time × drug interaction (*F* [1,65] = 4.37, *p* = 0.04, *η*_*p*_^*2*^ = .06). Post-hoc paired samples t-tests showed that there was a decrease in craving from session one (pre-drug) to session two (post-drug) in both drug groups, but this effect was larger in the varenicline group (see Table 2). For cigarette craving VAS data, the main effects of time (*F* [1,61] = 56.62, *p* < .001, *η*_*p*_^*2*^ = .48) and drug (*F* [1,61] = 16.32, *p* < .001, *η*_*p*_^*2*^ = .21) were replicated but there were no other effects or interactions (*ps* > .34).

**Table 2:** Craving (QSU) scores from pre-to post drug administration in varenicline and placebo groups

|  | Mean difference<br>from session one<br>to session two<br>(SD) | Effect size<br>(dz) | 95% CI | p-value |
| --- | --- | --- | --- | --- |
| Placebo | -4.7 (10.3) | 0.46 | -8.4 to -1.1 | 0.013 |
| Varenicline | -10.1 (10.6) | 0.95 | -13.8 to -6.4 | <0.001 |

#### Mood (VAS) across drug treatment

There was evidence of effects of time for happiness (*F* [1,62] = 4.37, *p* = .041, *η*_*p*_^*2*^ = .07), drowsiness (*F* [1,62] = 3.78, *p* = .057, *η*_*p*_^*2*^ = .06), depression (*F* [1,62] = 9.93, *p* = .003, *η* ^*2*^ = .14), anxiety (*F* [1,62] = 3.22, *p* =.078, *η*_*p*_^*2*^ = .05) and irritability (*F* [1,62] = 8.56, *p* = .005 *η*_*p*_^*2*^ = .12), with decreases in happiness, and increases in drowsiness, depression, anxiety and irritability. There was also evidence of an effect of drug for anxiety (*F* [1,62] = 9.01, *p* = .004, *η*_*p*_^*2*^ = .13), with lower anxiety reported in the varenicline group. There was no clear evidence of any other main effects or interactions (*ps* >.10).

#### Withdrawal across CBM (pre-CBM to post-scan)

There was evidence of a main effect of time (*F* [1,62] = 14.98, *p* < .001, *η*_*p*_^*2*^ = .20) with increases in withdrawal pre-CBM (*M =* 8.6, *SD* = 7.3) to post-CBM (*M =* 11.3, *SD* = 7.7). There was no clear evidence for other main effects or interactions (*ps* > .14).

#### Craving (QSU) across CBM (pre-CBM to post-scan)

For QSU data, there was evidence of effects of time (*F* [1,62] = 62.6, *p* < .001, *η*_*p*_^*2*^ = .50) and drug (*F* [1,62] = 8.82, *p* = .004, *η* ^*2*^ = .13), indicating increases in craving from pre-CBM (*M =* 27.7, *SD* = 12.1) to post scan (*M =* 38.5, *SD* = 12.8) and higher craving in the placebo group (*M =* 36.9, *SD* = 9.7) compared to varenicline group (*M =* 29.3, *SD* = 11.6). There was no strong evidence of other main effects or interactions (*p*s > .08). The main effects of time (*F* [1,62] = 56.62, *p* < .001, *η*_*p*_^*2*^ = .48) and drug (*F* [1,62] = 16.32, *p* < .001, *η*_*p*_^*2*^ = .21) were replicated using craving VAS data, and there was no clear evidence of any other effects or interactions (*ps* >.34).

#### Mood across CBM session (pre-CBM to post-scan)

There was evidence of main effects of time for happiness (*F*(1,61) = 4.24, *p* = .044, *η*_*p*_^*2*^ = .07), drowsiness (*F*(1,61) = 12.86, *p* = .001, *η*_*p*_^*2*^ = .17), energy (*F*(1,61) = 8.24, *p* = .006, *η*_*p*_^*2*^ = .12) and irritability (*F*(1,61) = 7.71, *p* = .007, *η*_*p*_^*2*^ = .11), with decreases in happiness and energy, and increases in drowsiness and irritability across the session. There was evidence of a main effect of drug (*F*(1,61) = 6.46, *p* = .014, *η*_*p*_^*2*^ = .10) and a time × drug interaction (*F*(1,61) = 6.15, *p* = .016, *η*_*p*_^*2*^ = .09) for anxiety, with higher anxiety in the placebo group (*M* = 25.4, *SD* = 19.9) compared to varenicline group (*M* = 15.1, *SD* = 14.0). Post-hoc paired t-tests indicated a decrease in anxiety across the session in the placebo group (*t* = 2.35, *df* = 33, *p* = .025, *dz =* .40), but not the varenicline group (*t* = -1.25, *df* = 32, *p* = .22, *dz =* .22). Finally there was evidence of a drug × CBM interaction for happiness (*F*(2,61) = 4.36, *p* = .017, *η*_*p*_^*2*^ = .13), with the varenicline group reporting lower happiness than the placebo group but only in the attend CBM condition.

### Neural response (fMRI data)

Following pre-processing, three participants (2 in the varenicline-attend group, 1 in the control-placebo group) were excluded from fMRI imaging analysis (2 due to excessive motion and 1 due to poor signal quality). There was no evidence of main effects of drug or CBM within the *a priori* mask. There was strong evidence of a drug × CBM interaction in one cluster (k = 32) in the right middle temporal gyrus (peak voxel: x = 58, y = −58, z = 12; z = 3.78, *p* <0.001). The placebo group exhibited greater temporal gyrus activation to smoking cues than the varenicline group in the control CBM condition (t = 2.86, *p* = .006) but less activation than the varenicline group in the toward CBM condition (t = 2.41, *p* = .02) (Figure 1).

**Figure 1:**
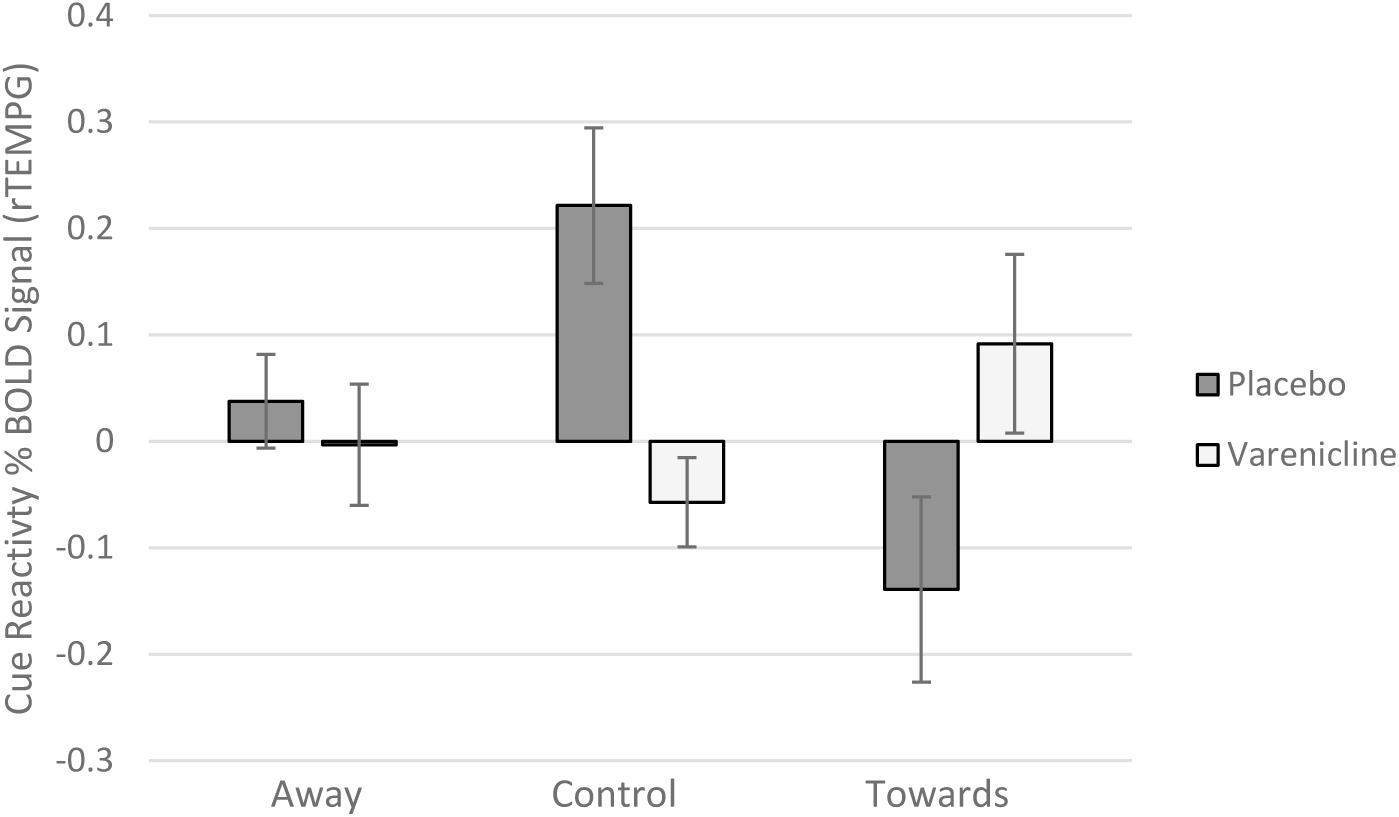
Neural response to smoking cues (% BOLD signal) in the right temporal gyrus across the three CBM groups following 1-week administration of placebo or varenicline.

Cognitive bias data from the visual dot probe task used in the regression analysis were transformed as described above. One subject with useable imaging data was excluded from the analysis due to incomplete behavioural data. Correlation between post-training bias scores and smoking cue reactivity (i.e., smoking greater than control cue contrasts) were put into a regression analysis. There was no clear evidence for areas of activation (*ps* >0.001).

## Discussion

We investigated the effects of CBM and one-week varenicline treatment on neural response to smoking-related cues in current smokers. There was no evidence that CBM training alone altered cognitive bias (post-CBM visual dot probe, Stroop), mood, craving or neural response to smoking-cues. There was evidence of reduced craving in the varenicline group as evinced by several drug by time interactions. Across the drug administration phase of the study (i.e., the six days preceding the study session), there was a greater reduction in QSU scores for the varenicline compared to placebo group. A main effect of drug was also evident for VAS craving scores (i.e., lower craving in varenicline group); however, the drug by time interaction was not replicated. The lower reports of craving in the varenicline group were also evident at the study session, during which CBM was administered. Finally, there was weak evidence that varenicline may have attenuated CBM-induced smoking bias, as there was an increase in smoking bias following CBM attend training, but only in the placebo group. There was no evidence of an interaction between varenicline and CBM on withdrawal, craving or mood.

There was evidence of a drug by CBM interaction on neural responses in one region within our *a priori* cue-reactivity mask. In right middle temporal gyrus (rMTG), activation in response to smoking relative to neutral cues was greater in the placebo as compared to the varenicline condition in the control CBM condition; the opposite pattern was observed in the toward CBM condition. Whereas the rTMG has been shown to be active in response to viewing smoking cues in meta-analyses (Tang et al., 2012), little has been reported regarding its potential role in processing conditioned smoking cues. Activation in the specific rTMG location we observed, has also been observed in studies of Theory of Mind, or the social-cognitive ability to infer the emotional and motivational experience (Vollm et al., 2006), where it may be involved in retrieval of memories associated with the behaviour of others. In the context of the present study, varenicline in the control CBM condition may decrease the degree to which such memories are accessed due to decreased tonic craving. Why toward CBM training would reverse such effects is unclear from the data, which suggests that additional research is needed to fully understand the influence of CBM on the processing of smoking cues.

Taken together these findings support a benefit of varenicline on tonic craving and neural response to smoking cues (which may be driven by the craving effects). While the effects of varenicline may be small, they are meaningful given the fact that the dosing regime delivered in the study is substantially lower than the clinically prescribed dose (i.e., 1 week compared to a standard 12-week course). However, we found no evidence of a benefit of CBM on any outcomes, and little evidence that varenicline would be a useful adjunct to smoking-related CBM. The CBM by drug interaction that was observed for the fMRI data, indicated that the effects of varenicline may have been attenuated for active CBM (i.e., the effects were only observed in the control training group). However, numbers are small and therefore this effect requires replication.

It is noteworthy that we did not find effects of CBM on measures of cognitive bias (visual dot probe and Stroop). There are known issues with the reliability of cognitive bias tests (Ataya et al., 2012), and therefore this may be a failure of the measure rather than a lack of effect. However, this indicates that the CBM may not have been effective, and these findings should be interpreted with this in mind. We hypothesised that effects of CBM would be potentiated by varenicline and our failure to observe such effects may be due to there being no CBM effects to strengthen. It is plausible that varenicline may potential effects of CBM if these effects can be reliably achieved.

There are some limitations of this study that should be considered when interpreting these findings. First, our sample size was small for the analysis of interactions. Our planned recruitment of 72 participants was achieved but not all participants were tested to completion, and our final sample was lower (n = 67 for subjective and cognitive data; n = 64 for fMRI data). We also have a computer malfunction for one of the conditions that was not identified until data were extracted. We had to replace a number of participants in one CBM condition (avoid) and therefore these individuals were tested outside of the randomisation sequence. We do not however expect that this had a substantial effect on outcomes as these individuals were testing in close time proximity to the rest of the sample. Furthermore, the researchers collecting data were not aware of the reason for additional recruitment, and therefore double-blinding was maintained. Third, our study recruited non-treatment seeking smokers, and it is plausible that effects of CBM may be stronger in individuals seeking treatment.

This study investigated neural responses to smoking cues following varenicline and CBM treatment. There was little evidence of neural effects of either drug or CBM. However, there was evidence of reductions in craving among smokers who completed one-week of varenicline treatment. Drug by CBM interactions were exploratory due to small sample sizes, but we observed an interaction on right temporal gyrus activity. Specifically, varenicline appeared to attenuate cue-related activity in the right temporal gyrus that was presented in the placebo group. However, this effect should be replicated in future research. In summary, this study finds little evidence of clinical potential of CBM.

## Abbreviations

CBM: Cognitive bias modification
PEQ-R: Eysenck Personality Questionnaire-Revised
MNWS: Minnesota Nicotine Withdrawal Scale
RT: reaction time
VAS: visual analogue scales

## Author contributions

MRM, FJM, SA and ASA contributed to the conception and design of the trial, and plans for data analysis. SA and ASA participated in data collection and project management. FJM and RK analysed and interpreted fMRI data. TW leads the clinical team (including DC, AG and KS) for subject recruitment. ASA drafted the manuscript, and all authors discussed, read and revised the manuscript. All authors approved the publication of this protocol.

## Acknowledgements

This study was supported by a Pfizer Ltd. Investigator Initiated Grant (grant ID: WS676950). MRM, ASA and SA are members of the UK Centre for Tobacco and Alcohol Studies, a UKCRC Public Health Research Centre of Excellence. ASA and MRM are members of the MRC Integrative Epidemiology Unit at the University of Bristol. We thank Aileen Wilson, Emma Howell, Alexander Davidson, George Stothart and Olivia Maynard for their assistance with data collection.

